# SimNIBS 2.1: A Comprehensive Pipeline for Individualized Electric Field Modelling for Transcranial Brain Stimulation

**DOI:** 10.1101/500314

**Authors:** Guilherme B Saturnino, Oula Puonti, Jesper D Nielsen, Daria Antonenko, Kristoffer H Madsen, Axel Thielscher

## Abstract

Numerical simulation of the electric fields induced by Non-Invasive Brain Stimulation (NIBS), using realistic anatomical head models has gained interest in recent years for understanding the NIBS effects in individual subjects. Although automated tools for generating the head models and performing the electric field simulations have become available, individualized modelling is still not standard practice in NIBS studies. This is likely partly explained by the lack of robustness and usability of the previously available software tools, and partly by the still developing understanding of the link between physiological effects and electric field distributions in the brain. To facilitate individualized modelling in NIBS, we have introduced the SimNIBS (Simulation of NIBS) software package, providing easy-to-use automated tools for electric field modelling. In this article, we give an overview of the modelling pipeline in SimNIBS 2.1, with step-by-step examples of how to run a simulation. Furthermore, we demonstrate a set of scripts for extracting average electric fields for a group of subjects, and finally demonstrate the accuracy of automated placement of standard electrode montages on the head model. SimNIBS 2.1 is freely available at www.simnibs.org.

## 1. Introduction

Non-Invasive Brain Stimulation (NIBS) aims at modulating brain activity by inducing electric fields in the brain [1]. The electric fields are generated either by a magnetic coil, in case of transcranial magnetic stimulation (TMS), or by a current source and electrodes placed directly on the scalp, in case of transcranial electric stimulation (TES). In both cases, the induced electric fields in the brain have a complex and often counter-intuitive spatial distribution, which is dependent on the individual anatomy of a target subject. In recent years, there has been a growing interest in moving away from a one-size-fits-all stimulation approach in NIBS towards more individually informed protocols [2]. The driving force behind this shift is the widely reported variation of NIBS effects within and between individuals [3], which could be explained in part by the interplay of the individual anatomy and the electric field propagation [4]. Although software tools have become available that generate realistic anatomical models of the head based on magnetic resonance imaging (MRI) scans and use those models to numerically estimate the electric field induced in the brain, they are still not standardly used in NIBS studies. This is likely also due to the lack of robustness and usability of the previous generation of tools, in turn hampering the individualized application of NIBS in both mapping the human brain function and as a rehabilitation tool in various neuropathologies [5], [6].

The aim of SimNIBS is to facilitate the use of individualized stimulation modelling by providing easy-to-use software tools for creating head models, setting up electric field simulations, and visualizing and post-processing the results both at individual and group levels. SimNIBS was first released in 2013 [7], had a major update in 2015, with the release of version 2 [2], and more recently another major update with the release of version 2.1, described in the current work. SimNIBS 2.1 is free software, distributed under a GPL 3 license, and runs on all major operating systems (Windows, Linux and MacOS). In this tutorial, we will concentrate on **what** SimNIBS 2.1 can be used for and **how** the analyses are run in practice with step-by-step examples. The chapter is structured as follows: First, we give a general overview of the simulation pipeline and of its building blocks. Next, we provide a step-by-step example of how to run a simulation in a single subject, and then we demonstrate a set of MATLAB tools developed for easy processing of multiple subjects. Finally, we conclude with an analysis of the accuracy of automated electrode positioning approaches. More information, as well as detailed tutorials and documentation can be found from the website www.simnibs.org.

## 2. Overview of the SimNIBS workflow

Figure 1 shows an overview of the SimNIBS workflow for an individualized electric field simulation. The workflow starts with the subject’s anatomical MRI images, and optionally diffusion weighted MRI images. These images are segmented into major head tissues (white and grey matter, cerebrospinal fluid, skull and scalp). From the segmentations, a volume conductor model is created, and used for performing the electric field simulations. The simulations can be set-up in a graphical user interface (GUI) or by scripting. Finally, the results can be mapped into standard spaces, such as the MNI space or FreeSurfer’s FsAverage.

**Figure 1:**
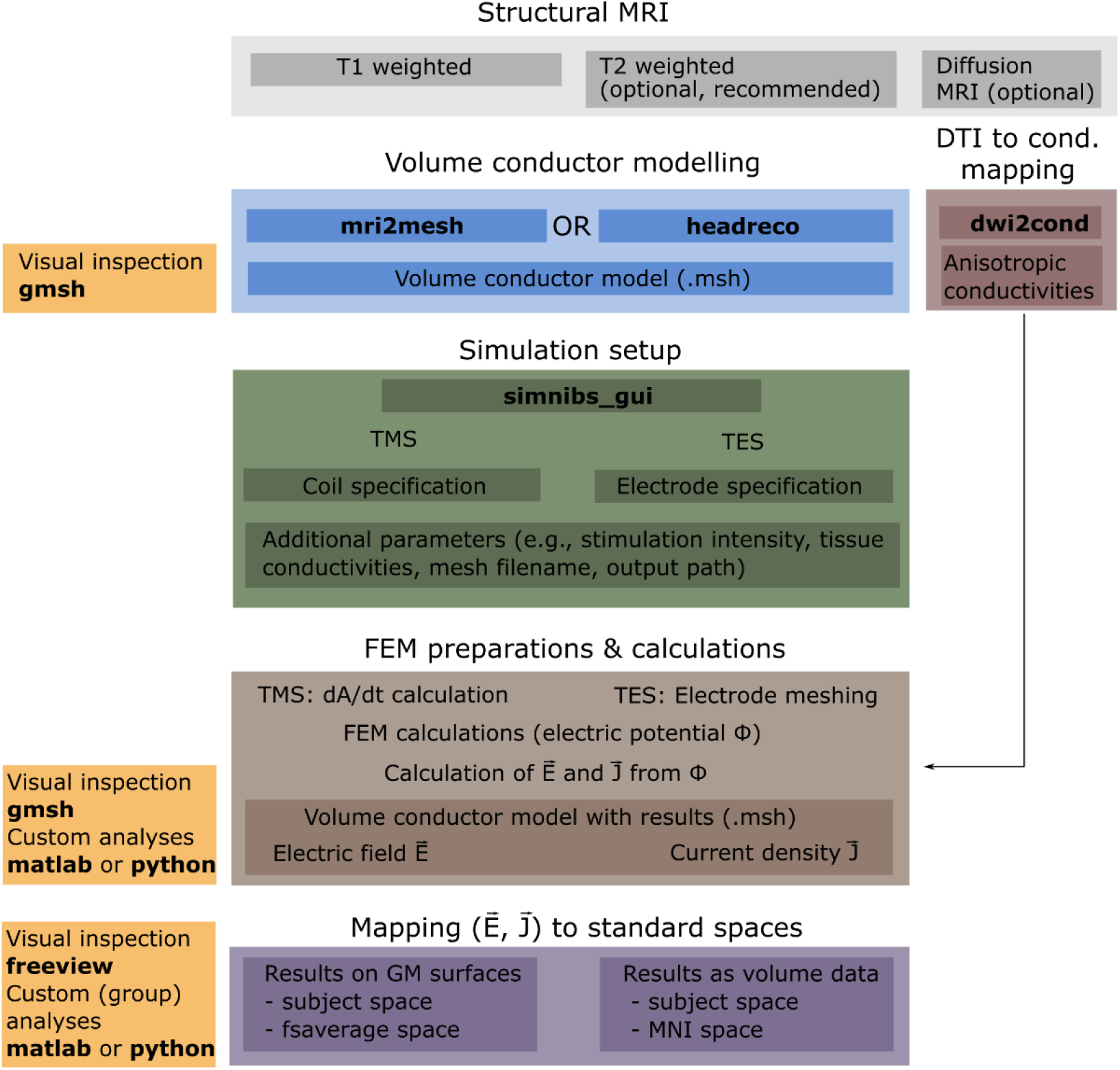
Overview of the SimNIBS workflow.

### 2.1 Structural magnetic resonance imaging scans

The minimum requirement for running an individualized SimNIBS simulation is a T1-weighted structural scan of a subject’s head anatomy. Although SimNIBS will run on almost all types of T1-weighted scans, we have found that setting the readout bandwidth low to ensure good signal-to-noise ratio in the brain region and using a fat suppression method, such as selective water excitation, to minimize the signal from spongy bone, typically ensure a high quality of the resulting head models. See Figure 2 for an example of good quality scans we found to work well with SimNIBS and [8] for the details of the sequences.

**Figure 2:**
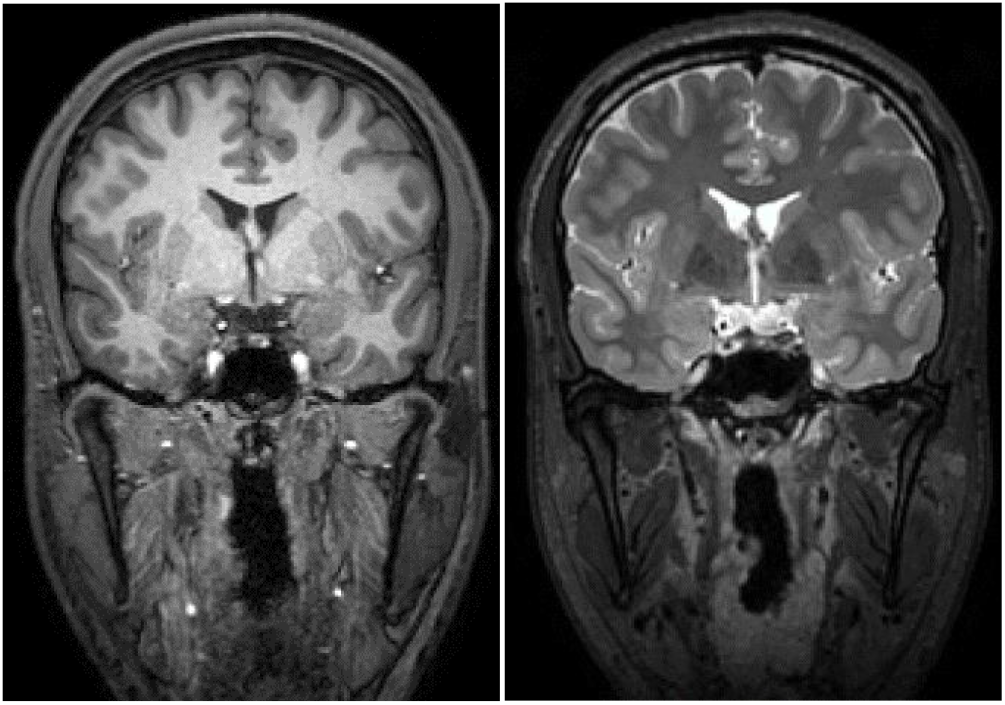
Example of high-quality T1- and T2-weighted scans likely to work well with SimNIBS. Note that in the T1-weighted scan the skull appears dark due to the fat-suppression.

Including a T2-weighted scan is optional, but highly recommended as it facilitates accurate segmentation of the border between skull and cerebrospinal fluid (CSF). Both skull and CSF appear dark in T1-weighted scans, whereas in T2-weighted scans the CSF lights up, thus guiding the separation between the tissues. Skull has a low electric conductivity while CSF is highly conducting, meaning that any segmentation errors in these two compartments can have a large effect on the resulting electric field distribution inside the head, especially when TES is applied [8]. If you are interested in modelling the neck region in detail, we recommend using neck coils if these are available at the imaging site.

Optionally, SimNIBS also supports modelling of anisotropic conductivities for grey (GM) and white matter (WM), which requires a diffusion weighted MRI scan (dMRI). Only single shell data (i.e., with a single b-value in addition to some b=0 images) with a single phase encoding direction for the echo planar imaging (EPI) readout is supported.

### 2.2 Volume Conductor Modelling

The first step in the pipeline is the generation of a volume conductor model of the head, which is needed for simulating the induced electric fields. In order to create this finite element (FEM) mesh, we need to assign each voxel in the MRI scan(s) to a specific tissue class, i.e., to segment the scan into the different head tissues. Currently, SimNIBS offers two options for segmentation: **mri2mesh** [7] and **headreco** [8].

**mri2mesh** combines FSL [9] (version 5.0.5 or newer) and FreeSurfer [10] (version 5.3.0 or newer) to segment the head tissues. FSL is used to segment the extra-cerebral tissues, while FreeSurfer is used to segment the brain and to generate accurate surface reconstructions of the grey matter sheet. Note that **mri2mesh** is restricted only to the head and does not create models of the neck region.

**headreco** uses the SPM12 [11] toolbox for segmenting the MRI scan, and is now the recommended option in SimNIBS. It has been shown to be more accurate in segmenting the extra-cerebral structures, especially the skull, compared to **mri2mesh** [8], while also providing accurate segmentations of the brain tissues. The computational anatomy toolbox (CAT12, recommended) [12] provided with SPM can be used to create surface reconstructions of the grey matter sheet which are on par with the accuracy of those generated by FreeSurfer [12]. In addition, **headreco** has an extended field-of-view, also modelling the neck region. For ease-of-use, both SPM12 and CAT12 are distributed together with SimNIBS.

Once the segmentation by either method has finished successfully, the tissue maps are cleaned by applying simple morphological operations, and used to create surface reconstructions. As a final step, the FEM mesh is generated by filling in tetrahedrons between the tissue surfaces using Gmsh [13].

Neither **mri2mesh** nor **headreco** have off-the-shelf support for pathologies such as tumours or lesions. These can however be included into the head models by manually editing the segmentation masks generated by the methods. When using **mri2mesh** please consult the FreeSurfer website (https://surfer.nmr.mgh.harvard.edu/fswiki/FsTutorial/WhiteMatterEdits_freeview) on how to handle scans with pathologies. Manual edits using **headreco** should be done on the output segmentation masks in the *mask_prep* folder located within the *m2m_{subID}* folder. Once corrections have been made, the surface meshing step (“**headreco surfacemesh subID**”) and volume meshing step (“**headreco volumemesh subID**”) should be re-run to generate the edited head model. Note that when creating head models from scans with pathologies the CAT12 toolbox should <u>not</u> be used.

**dwi2cond (optional)** uses FSL (version 5.0.5 or newer) to prepare diffusion tensors for GM and WM from dMRI data. The tensors are used by SimNIBS to estimate anisotropic conductivities in WM and GM during the FEM calculations.

### 2.3 Simulation setup

Simulations can be set-up using the graphical user interface (GUI), which provides an interactive view of the head model. This allows users to easily select parameters such as coil positions, electrode positions and shapes, as well as more advanced settings such as tissue conductivities and post-processing options.

It might also be of interest to do simulations of one or a few different set-ups across a group of subjects. With this in mind, version 2.1.1 introduced a new interface for setting up simulations using MATLAB or Python scripts.

The GUI as well as the scripts will be described in more detail in Section 3, as well as on the website www.simnibs.org.

### 2.4 Finite element method calculations

Transcranial direct current (tDCS) simulations begin by adding electrodes to the head model. In this step, nodes in the skin surface are shifted to form the shape of the electrode, while keeping good quality elements. Afterwards, the body of the electrodes is constructed by filling in tetrahedra. As this step does not require re-meshing the entire head, it can be done much more efficiently compared to other methods that require re-meshing, especially when only a few electrodes are used.

TMS simulations start by calculating the change in the magnetic vector potential ***A***, that is, *d**A***/*dt* the field in the elements of the volume conductor mesh for the appropriate coil model, position and current. There are currently two types of coil models:

**.ccd files**: Created from geometric models of the coil and represented as a set of magnetic dipoles from which we can calculate the *d**A***/*dt* field using a simple formula [14].

**.nii files**: Created either from geometric models of the coils or direct measurement of the magnetic field [15]. Here, the *d**A***/*dt* field is defined over a large volume, and the calculation of the *d**A***/*dt* at the mesh elements is done via interpolation. This allows for faster simulation set-up at little to no cost in simulation accuracy.

Both simulation problems are solved using the FEM with linear basis functions. This consists of constructing and solving a linear system of the type ***Mu*** = ***b***, where ***M*** is a large (in SimNIBS typically ~10^6^ × 10^6^) but sparse matrix, called “stiffness matrix”, ***u*** are the electric potentials at the nodes and the right-hand side ***b*** contains information about boundary conditions (such as potentials in electrode surfaces in tDCS simulations), and source terms (such as the *d**A***/*dt* field in TMS simulations). SimNIBS solves the linear system using an iterative preconditioned conjugate gradient method [16]. SimNIBS 2.1 uses GetDP [17] to form the linear system, which in turn calls PETSc [18] to solve it.

TDCS simulations can also be easily extended to simulations of transcranial alternating current stimulation (tACS). In the frequency ranges used in tACS, a quasi-static approximation holds [19]. In the quasi-static approximation, the relationship between input currents *I(t)* and the electric field at the positions ***x***, *E*(***x***) is linear:

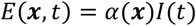

Where *α*(***x***) is a proportionality constant, meaning that it does not vary during the oscillation. This constant can be obtained simply by running a simulation where we set the input current to unity. *I*(*t*) is the input current. For example, a sinusoidal current input can be written as

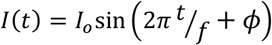

Where *f* is the stimulator frequency, *ϕ* the stimulator phase, and *I_o_* the stimulator amplitude, which corresponds to half of the peak-to-peak current. Usually, we would visualize the electric field at the maximum or minimum of *I*(*t*), which corresponds to ±*I_o_*. In case several stimulators are used at different frequencies of phases, we have several pairs (*α_i_*(***x***)*I_i_*(*t*)), one for each stimulator, and the total electric field at a given time point is given by the sum of their individual contributions

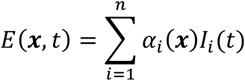

### 2.5 Mapping fields

The result of the FEM calculation is the electric field at each tetrahedral element of the subject’s head mesh. However, visualization is often easier using cortical surfaces or NifTI volumes. Therefore, SimNIBS 2.1 can transform fields from the native mesh format to these formats via interpolation. Our interpolation algorithm is based on the superconvergent patch recovery method [20], which ensures interpolated electric field values that are consistent with tissue boundaries.

When performing simulations on multiple subjects, we often want to be able to directly compare the electric field across subjects to, for example, correlate the electric field with behavioural or physiological data on the stimulation effects [21]. For this purpose, SimNIBS can also transform simulation results to the MNI template, using linear and non-linear co-registrations, as well as to the FreeSurfer’s FsAverage surface.

## 3 Practical examples and use cases

### 3.1 Hello SimNIBS: how to process a single subject

Here we describe how to run a TMS and a tDCS simulation on a single example subject. The example subject “Ernie” can be downloaded from the SimNIBS website, and the steps below can be reproduced step-by-step to get familiar with SimNIBS.

#### 3.1.1 Generating the volume conductor model

Open a terminal and go to the directory “ernie” to access the example data set. Copy the content of the “org”-subfolder to another location in order to not overwrite the files of the original example dataset. Next, go to the folder where you copied the data, and call headreco to generate the volume conductor model:

> **headreco all --cat ernie ernie_T1.nii.gz ernie_T2.nii.gz**

In the command, the first argument, “**all**”, tells headreco to run all reconstruction steps including: segmentation, clean-up of tissue maps, surface meshing, and volume meshing. The second argument, “--**cat**” is a flag for using the CAT12 toolbox for accurate reconstruction of the cortical surface. The third argument, “**ernie**” is a subject identifier (subID), which is used to name generated folders, e.g., m2m_ernie, and output files, e.g., ernie.msh. The two final arguments are the paths to the T1- and T2-weighted structural scans.

A few extra input options are useful to know:

-**d no-conform** adding this option will prevent headreco from modifying, i.e., transforming and resampling, the original MRI scan. This might be desirable when a one-to-one correspondence between the head model coordinates and the neural navigation system coordinates is required.
-**v** <**density**> this option allows you to set the resolution, or vertex density (nodes per mm^2^), of the FEM mesh surfaces. By default, SimNIBS uses 0.5 nodes/mm^2^ as the <**density**> value.

In general, we recommend using the --**cat** option; however, the execution time will be longer compared to omitting the option. In addition, if you want to process scans with pathologies, you should not use CAT12, as the cortical reconstruction is not designed to work with pathologies.

After **headreco** has finished please check the quality of the headmodel by calling:

> **headreco check ernie**

If needed, open a new terminal for that and go into the folder in which you started headreco the first time. For our example case the subject identifier is “ernie”, but please replace this one with whichever subID was used in the first call to **headreco**. Note that we recommend that you have installed freeview (provided by FreeSurfer) to visualize the results. The **check** function displays two windows for inspecting the output. The first window shows the T1-weighted scan with the segmentation and structure borders overlaid (Figure 3, left). We recommend de-selecting the segmentation (**ernie_final_contr.nii**) in freeview, and checking that the segmentation borders follow the intensity gradients of different tissues (Figure 3, middle). Figure 4 shows the second freeview window, which displays the T1-weighted scan co-registered to the MNI template. We recommend checking if the T1-weighted scan overlaps well with the MNI template by de-selecting the T1-weighted scan (**T1fs_nu_nonlin_MNI.nii**) in freeview (Figure 4, right). Figure 5 shows an example of a segmentation error where the skull is erroneously labelled as skin. This can be seen in the front of the head, where the skin label protrudes into the skull. This example emphasizes the need for fat-suppressed data when only a T1-weighted scan is used. In the scan shown in Figure 5, spongy bone is bright with intensities comparable to those of scalp, causing the segmentation method to mistake it as extra-cerebral soft tissue. Small segmentation errors like this can be corrected by manually relabelling the segmentation masks in the “*mask_prep*” folder located in the *m2m_{subID}* folder, and re-running the surface and volume meshing steps. If you are not familiar with using freeview, please check the tutorial on the SimNIBS website (*http://www.simnibs.org/_media/docu2.1.1/tutorial_2.1.pdf*). If you do not have access to freeview, the visualizations will be displayed using SPM. However, these are very primitive and are not recommended for checking the output from **headreco**.

**Figure 3:**
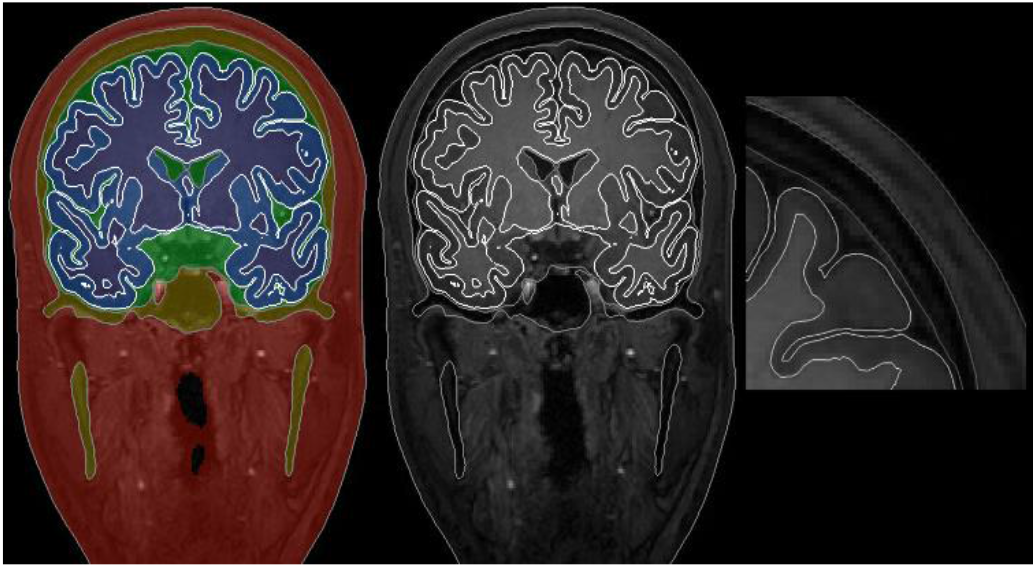
Data displayed after calling the **check** option. Left: T1-weighted scan with the segmentation and structure borders overlaid. Middle: structure borders overlaid on the T1-weighted scan after de-selecting the segmentation in freeview. Right: zoom-in of the cortex. Note that the segmentation borders nicely follow the intensity borders between the tissues.

**Figure 4:**
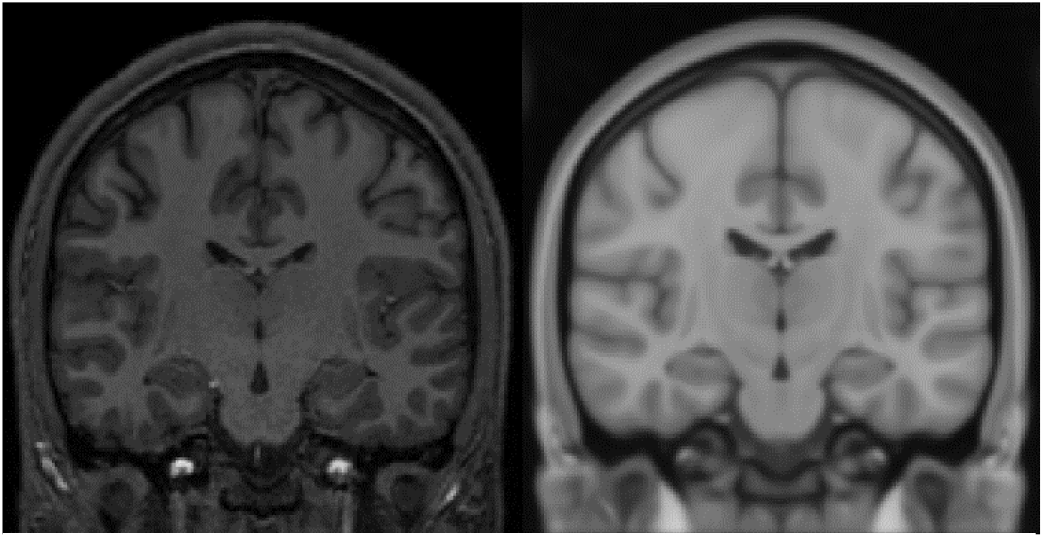
Data displayed after calling the **check** option. Left: T1-weighted scan co-registered to the MNI template. Right: MNI template shown after de-selecting the T1-weighted scan in freeview. Note that the scans seem to be well registered.

**Figure 5:**
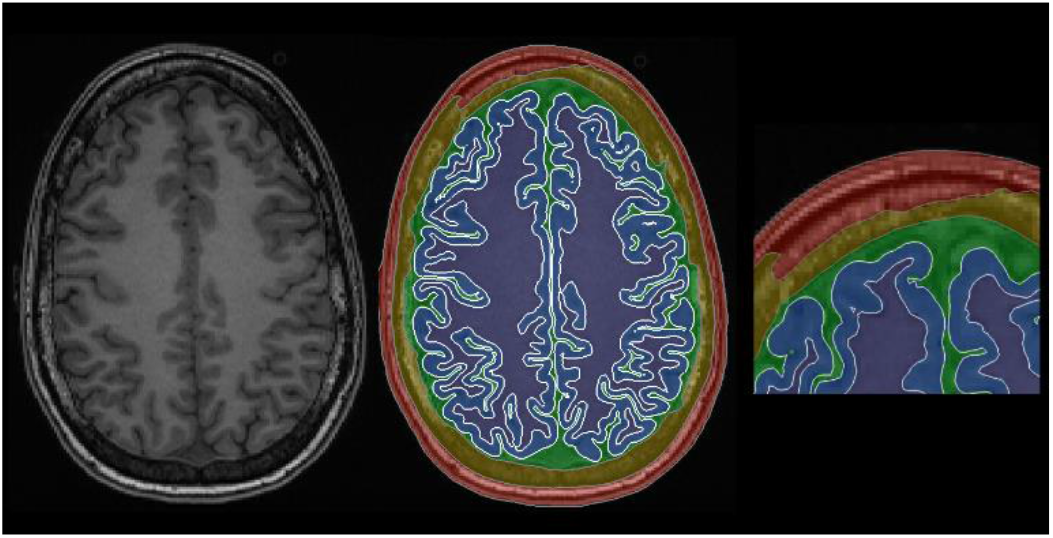
Example of a segmentation error after headreco processing. The spongy bone is erroneously labelled as skin. This example emphasizes the needfor fat-suppression when using only a T1-weighted scan.

Finally, you should inspect the volume conductor mesh for any obvious errors. This can be done by calling:

> **gmsh ernie.msh**

in the subject folder. The call opens a gmsh window displaying the generated headmodel, please see the tutorial on the website if you are not familiar with gmsh (*http://www.simnibs.org/_media/docu2.1.1/tutorial_2.1.pdf*).

The folder structure and most important files are shown in Table 1.

- *eeg_positions/* Folder containing the 10-10 electrode positions for the subject both as a “.*csv*”, used for acquiring electrode positions, and a “.geo” file, used for visualization of the positions in Gmsh. If you have custom electrode positions, they should be added here as a .csv file.
- *mask_prep/* Folder containing the cleaned tissue maps along with the white matter and pial surface files if CAT12 was used. In case there are errors in the segmentation, the masks can be manually corrected and a new head model can subsequently be generated. Note that the CAT12 WM and GM surfaces can currently not be modified.
- *headreco_log.html*, a log-file with output from the **headreco** run. If something goes wrong, the log-file helps with troubleshooting, and should be sent as an attachment when contacting the SimNIBS support email list.
- *ernie.msh*, the FEM head model used for the simulations.
- *ernie_T1fs_conform.nii.gz*, the input scan in the conform space defined by the –d option. This scan has the same millimetre space as the head model, and can be used to annotate landmarks which can then be directly transformed onto the head model.

**Table 1:**
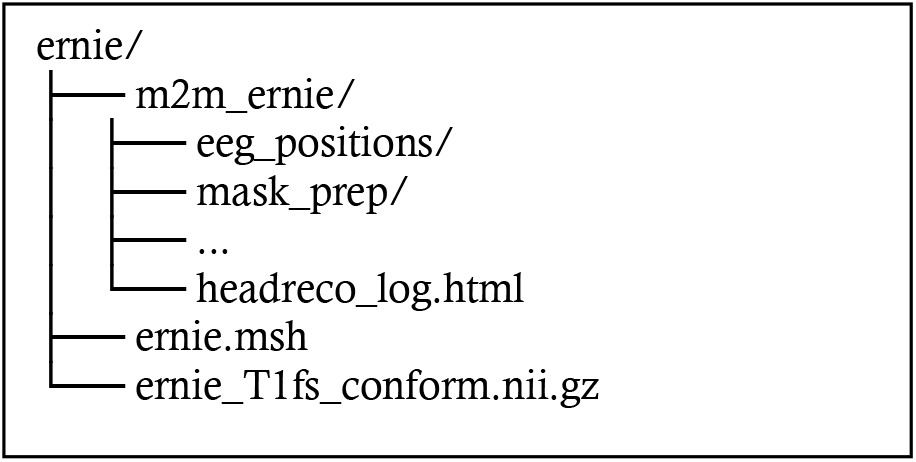
The folder structure after **headreco** has finished. In this table only the most important folders and files are listed.

#### 3.1.2 Setting up a simulation

Once the head model is ready, we can set-up tDCS and TMS simulations interactively using the GUI. The GUI can be started on the command line by calling:

> **simnibs_gui**

There, the user can:

- Visualize and interact with head models.
- Define electrode and coil positions by clicking in the model or selecting a position from the EEG 10-10 system.
- Visually define electrode shapes and sizes.
- Select from the available coil models.
- Change tissue conductivity parameters and set-up simulations with anisotropic conductivity distributions.
- Run simulations.

In the GUI, there are two types of tabs, one for tDCS simulations, and another for TMS simulations, shown respectively in the top and bottom of Figure 6. TDCS tabs define a single tDCS field simulation with an arbitrary number of electrodes. On the other hand, TMS tabs can define several TMS field simulations using the same coil. For this example, we will set-up a tDCS simulation with a 5×5 cm anode placed over C3 and a 7×5 cm cathode placed over AF4, and a TMS simulation with the coil placed over the motor cortex, pointing posteriorly. Details on how to use the graphical interface can be found on the website (*http://www.simnibs.org/_media/docu2.1.1/tutorial_2.1.pdf*).

**Figure 6:**
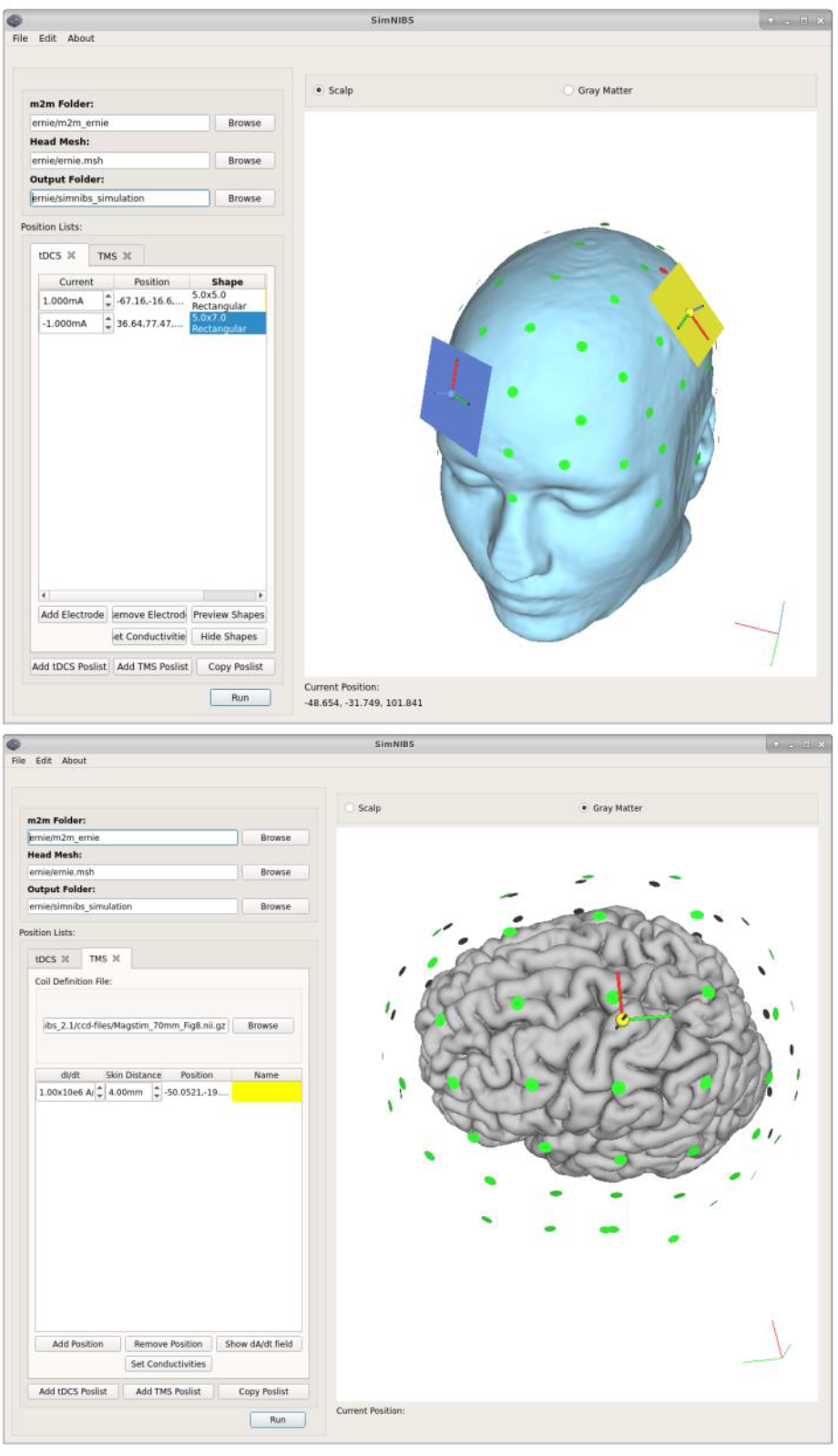
Set-up of a tDCS (top) and a TMS (bottom) simulation in the graphical user interface.

After the simulation set-up, click on the **Run** button to start the simulations. Running both simulations takes 10 to 15 minutes, depending on the computer, and uses around 6 GB of memory. As a note, before starting the simulations, you can set additional options (in the menu Edit➔Simulation Options) to let SimNIBS write out the results as surface data or NifTI volume data. This is not further covered in this basic example, but the output files created in these cases are described in the next example. The results of the simulation will be written in the output folder specified in the GUI, in this case “*simnibs_simulation/*”. The folder has the files shown below in Table 2.

- “*ernie_TDCS_1_scalar.msh*” is the output from the tDCS simulation, in Gmsh “.msh” format. The first part of the file name, “ernie”, is the subID. The second part, “*TDCS*”, informs us that this is a tDCS simulation. The third part, “*1*”, that this was the first simulation we have defined in the GUI, and finally, “*scalar*” tells us have used scalar (as opposed to anisotropic) conductivities for the simulations.
- “*ernie_TMS_2-0001_Magstim_70mm_Fig8_nii_scalar.msh*” is the output of the second simulation, also in gmsh “.*msh*” format. As is the case for the tDCS output, the first part of the file name is the subID, and the second is the number of the simulation in the simulation list. Afterwards, we see the number of the TMS position, as it might happen that several TMS positions are defined in a single TMS list. Afterwards, “*Magstim_40mm_Fig8_nii*” gives us the name of the coil used for the simulation, and “*scalar*” the type of conductivity.
- “*ernie_TMS_2-0001_Magstim70mm_Fig8nii_coil_pos.geo*” is a Gmsh “.geo” file which shows the coil position for the corresponding simulation.
- “*simnibs_simulation_20180920-13041.log*” is a text file with a detailed log of the simulation steps. This file can be used for troubleshooting. Here, the second part of the file is date and time information of when the simulation started.
- “*simnibs_simulation_20180920-13041.mat*” is a MATLAB data file with the simulation set-ups. This file can be loaded into the GUI or Matlab at later time points to check the simulation parameters, or to change them and re-run the simulation.

**Table 2:**
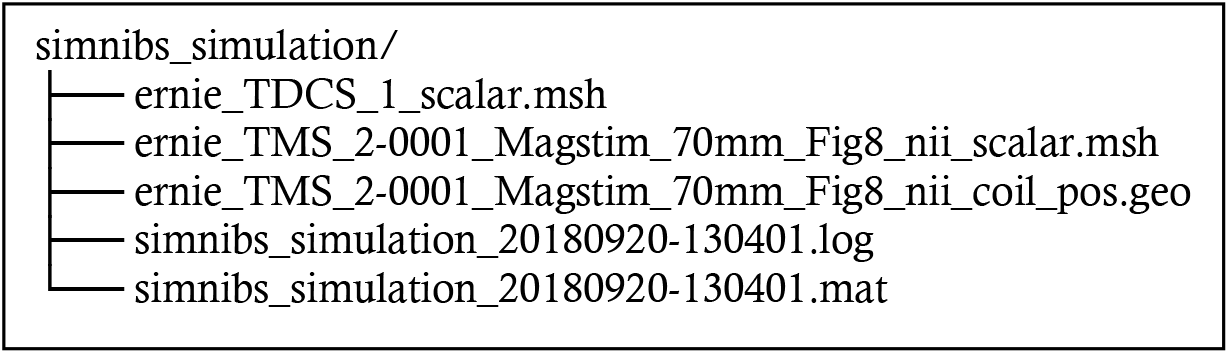
The outputfolder of a simple tDCS and TMS simulation.

#### 3.1.3 Visualizing fields

The electric field ***E*** is a vector field which means that the electric field has both a norm (i.e., vector length or magnitude) and a direction in space, as shown in Figure 7. As visualizations of the entire vector is challenging and often unclear, in SimNIBS we usually visualize the **norm** (or strength) of the electric field instead. The norm of the electric field corresponds to the size of the electric field vector, and therefore is always positive and does not contain any information about the direction of the electric field.

**Figure 7:**
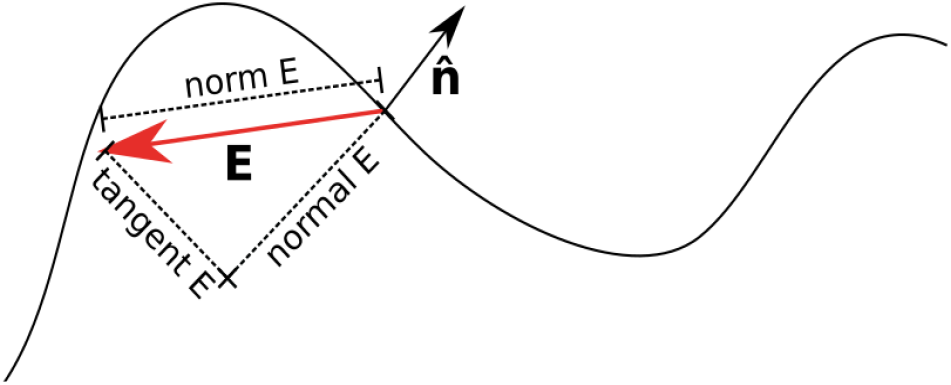
Decomposition of a vector **E** in relation to a surface. The norm corresponds to the length of the vector. At each point, the surface defines a normal vector 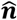, this vector is perpendicular to the tangent plane to the surface at that point. Given the normal vector, we can decompose the vector **E** in a normal and a tangent component. The normal component is the part of **E** in the same line as the normal vector, and the tangent component is perpendicular to it. The normal component also has a sign, indicating if the field is entering or leaving the surface. In SimNIBS, a positive normal indicate the field is entering the surface, and a negative normal indicate the field is leaving the surface.

One way we can quickly visualize the simulation results is to use the **mesh_show_results** MATLAB function. This function comes as a part of SimNIBS version 2.1.2, and provides visualizations of the output fields using MATLAB plotting tools, as well as some summary values for the field strength and focality. For example, when running the function on the output tDCS mesh, we obtain the plot shown in Figure 8A, and the values below in Table 3.

**Figure 8:**
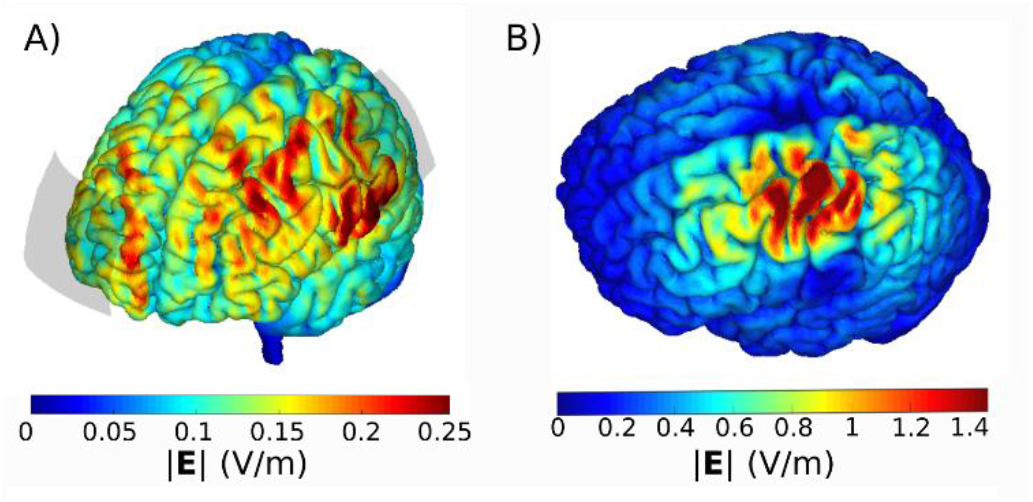
Visualization of A) tDCS and b) TMS electric field norms in MATLAB.

**Table 3:**
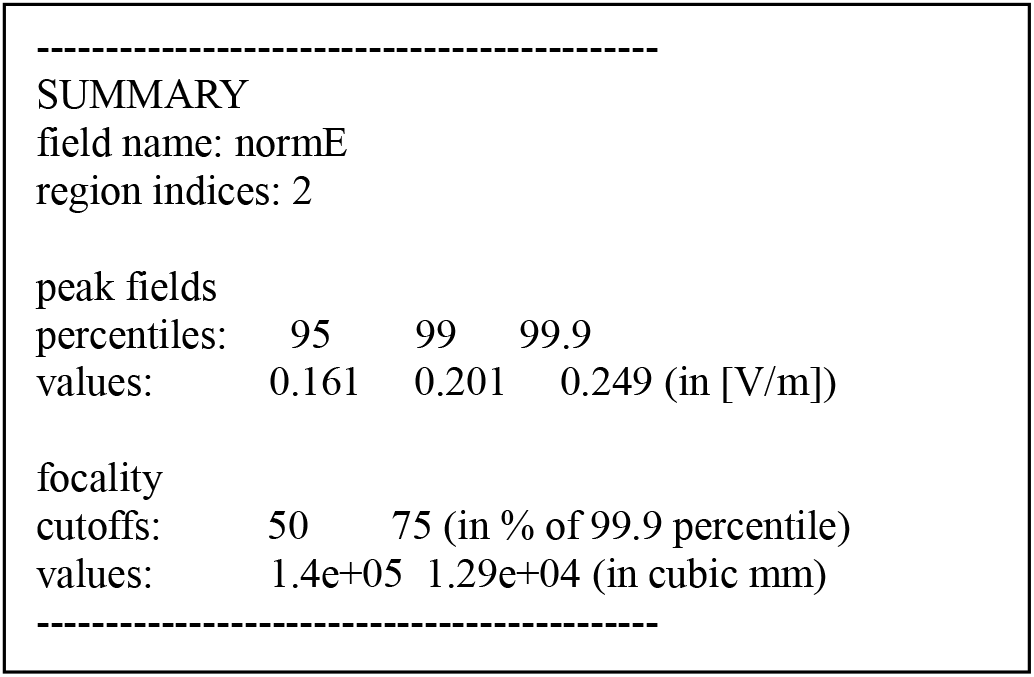
Output of **mesh_show_results** for the tDCS simulation.

peak fields
|  |  |  |  |
| --- | --- | --- | --- |
| percentiles: | 95 | 99 | 99.9 |
| values: | 0.161 | 0.201 | 0.249 (in [V/m]) |

|  |  |  |
| --- | --- | --- |
| cutoffs: | 50 | 75 (in % of 99.9 percentile) |
| values: | 1.4e+05 | 1.29e+04 (in cubic mm) |

The first lines in Table 3 show that the displayed data is the field “norm E”, that is, the norm or strength of the electric field, calculated in the region number 2, which corresponds to the GM volume. Afterwards, we have information on the peak electric fields. We see that the value of 0.161 V/m corresponds to the 95^th^ percentile of the norm of the electric field, the value of 0.201 V/m to the 99^th^ percentile and 0.249 to the 99.9^th^ percentile. We also have information about the focality of the electric field. Here, focality is measured as the GM volume with an electric field greater or equal to 50% or 75% of the peak value. To avoid the effect of outliers, the peak value is defined as the 99.9^th^ percentile.

Running the same function on the TMS result file, we obtain the plot shown in Fig. 8B, as well as the peak fields and focality measures shown below in Table 4.

**Table 4:**
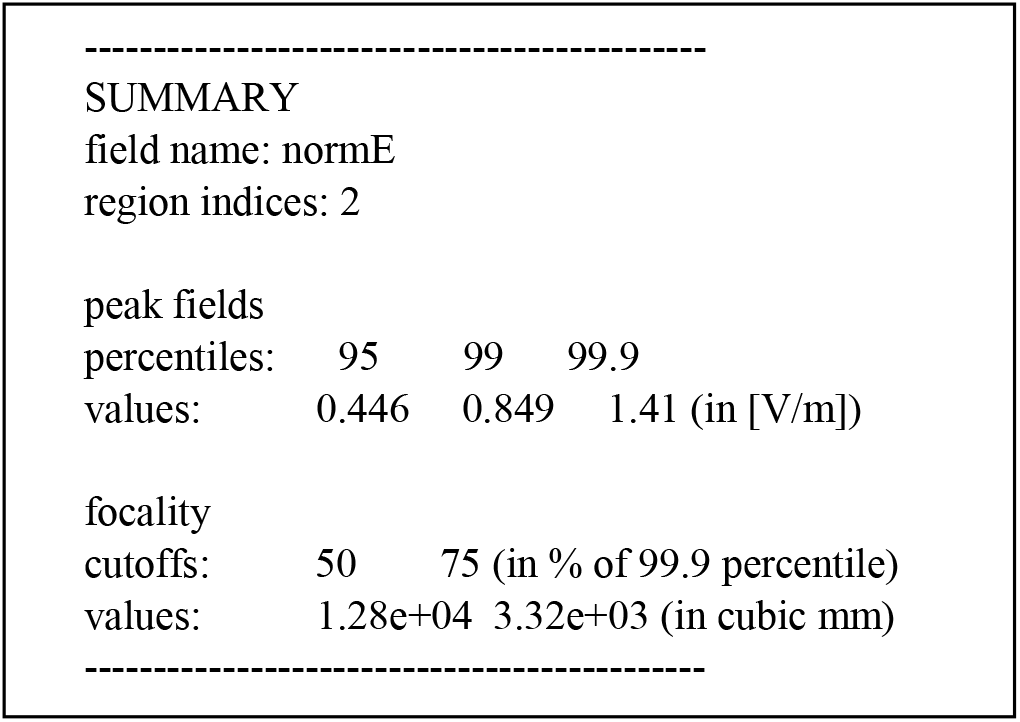
Output of **mesh_show_results** for the TMS simulation.

peak fields
|  |  |  |  |
| --- | --- | --- | --- |
| percentiles: | 95 | 99 | 99.9 |
| values: | 0.446 | 0.849 | 1.41 (in [V/m]) |

|  |  |  |
| --- | --- | --- |
| cutoffs: | 50 | 75 (in % of 99.9 percentile) |
| values: | 1.28e+04 | 3.32e+03 (in cubic mm) |

We can see that the peak fields for TMS are much higher than for tDCS, even though we simulated with a current of 10^6^ A/s, very low for TMS. In the focality measures, we see that the TMS electric fields are much more focal than the tDCS electric fields, with around five times less GM volume exceeding 75% of the peak value than tDCS.

Additionally, the “.*msh*” files can be opened with the Gmsh viewer, producing 3D visualizations as shown in Figure 9. Gmsh has a vast range of functionalities such as clipping planes, but can be harder to use than **mesh_show_results**.

**Figure 9:**
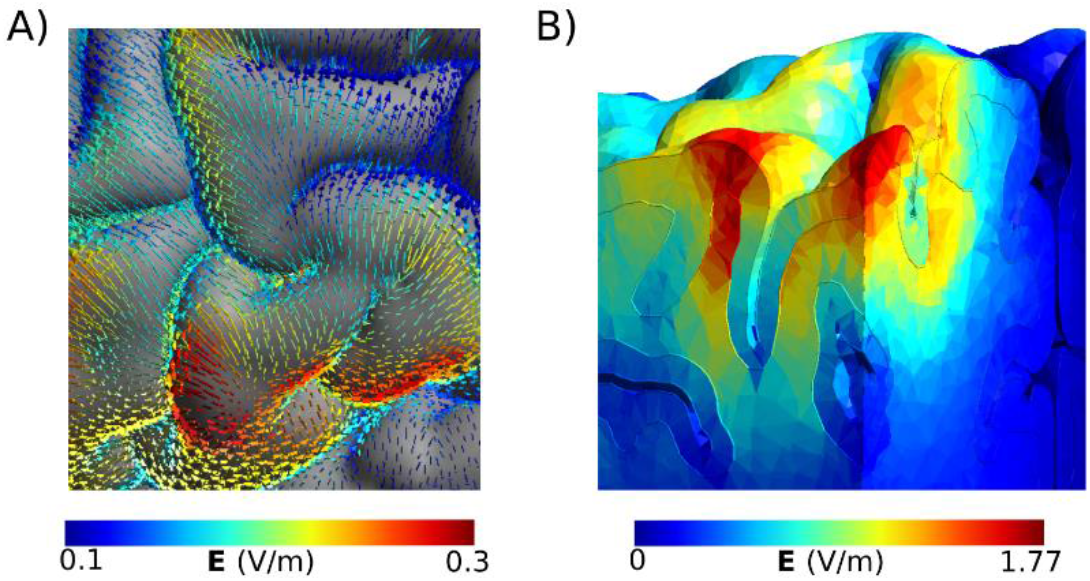
Visualization in Gmsh of A) electric field vectors around central gyrus for the tDCS simulation B) TMS electric field depth profile in the hotspot.

### 3.2 Advanced usage: group analysis

Now, we want to simulate one tDCS montage, with a 5×5 cm electrode over C3 and a 5×7 cm electrode over AF4 in five subjects, called “sub01”, “sub09”, “sub10”, “sub12”, “sub15” and visualize the results in a common space, namely the FsAverage surface. The subjects and example scripts can be downloaded from: https://osf.io/ah5eu/

#### 3.2.1 Head meshing

For each subject, follow the steps in section 3.1.1.

#### 3.2.3 Write a python or matlab script

We can set up the simulation of each subject using the GUI, as described in the first example. However, when working with multiple subjects, it can be advantageous to script the simulations for efficiency. SimNIBS provides both Matlab and Python interfaces to set up simulations. Script 1 shows how to set up and run a simulation with 5×5cm anode placed over C3 and a 7×5cm cathode placed over AF4 for all subjects. The output of Script 1 for sub01 is shown in Table 5.

**Script 1:**
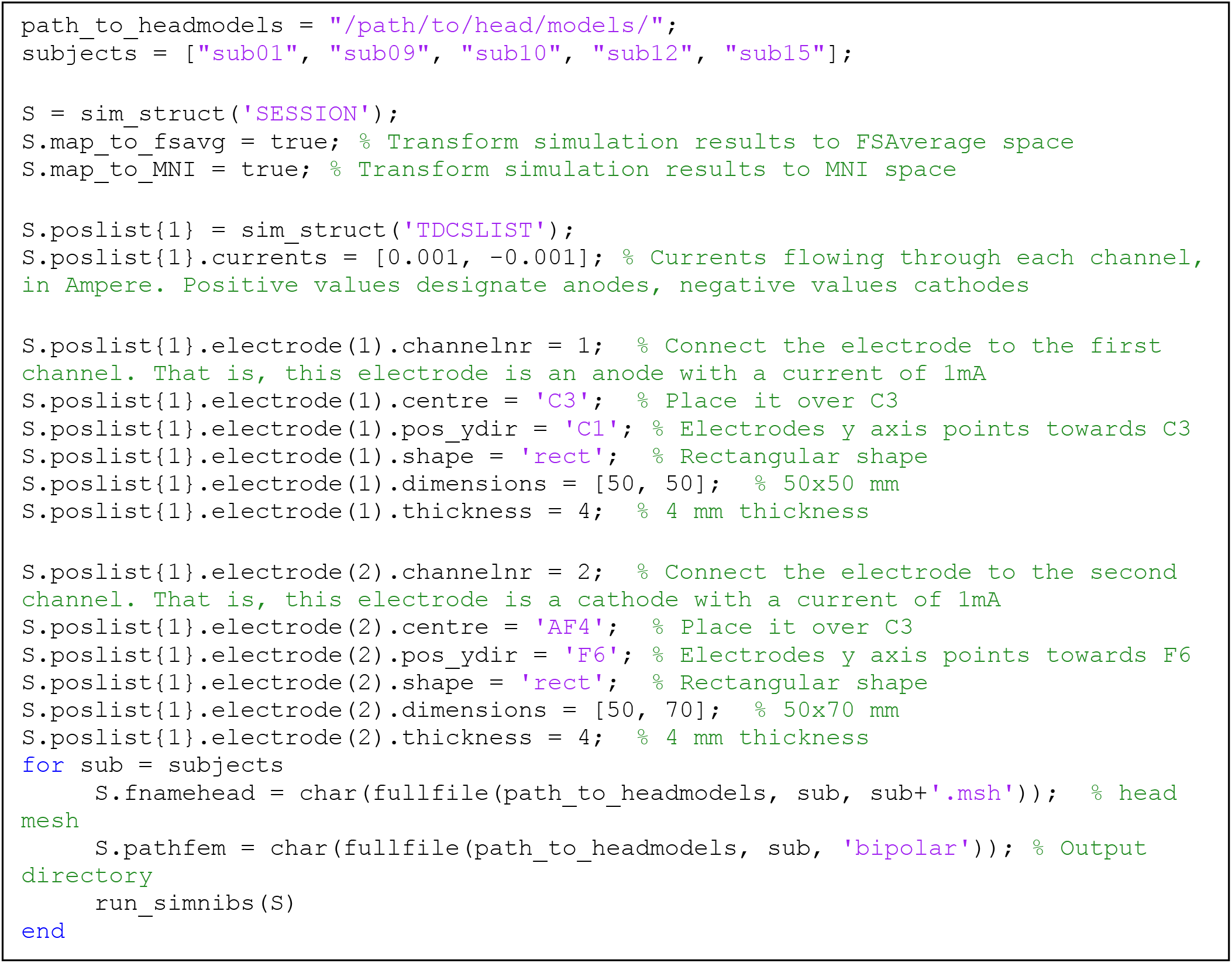
Script for running a tDCS simulation with an anode over C3 and a cathode over AF4 in five subjects and transforming the results to FSAverage and MNI spaces.

**Table 5:**
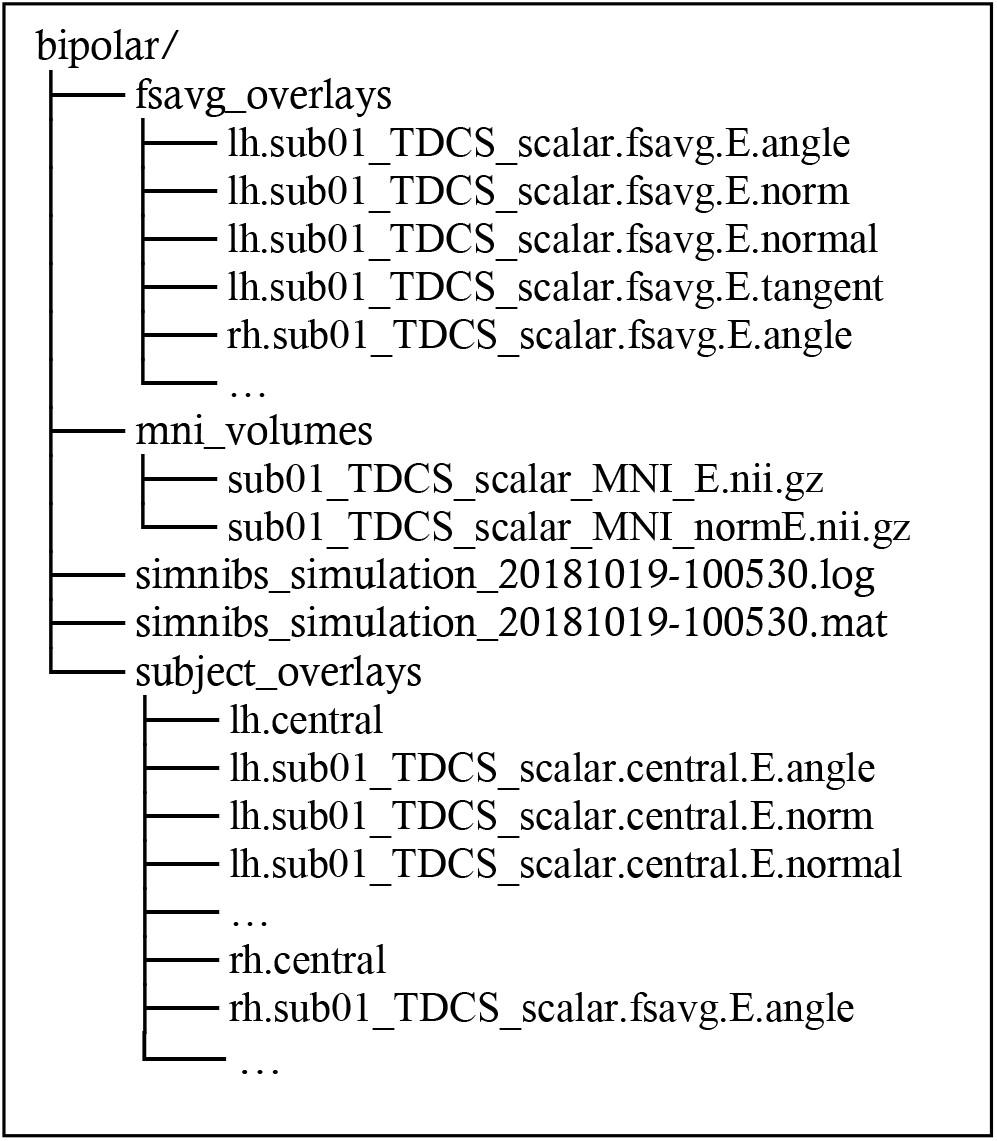
Output files and folders of Script 1 for sub01. The “.angle”,”.norm”,.. files are FreeSurfer overlay files and the “.central” files are FreeSurfer surface files

To define the rectangular electrodes, we need two coordinates. The “*centre*” defines where the electrode will be centred, and “*pos_ydir*” how the electrode will be rotated. More precisely, the electrode’s “y” axis is defined as a unit vector starting in “centre” and pointing towards “*pos_ydir*”. Figure 10 shows one of the cathodes (return electrode) defined using the script above, with the coordinate system and EEG positions overlaid. We can see that the electrode is centered in AF4, and its Y axis points towards F6. “*pos_ydir*” does not need to be set when the electrodes are round.

**Figure 10:**
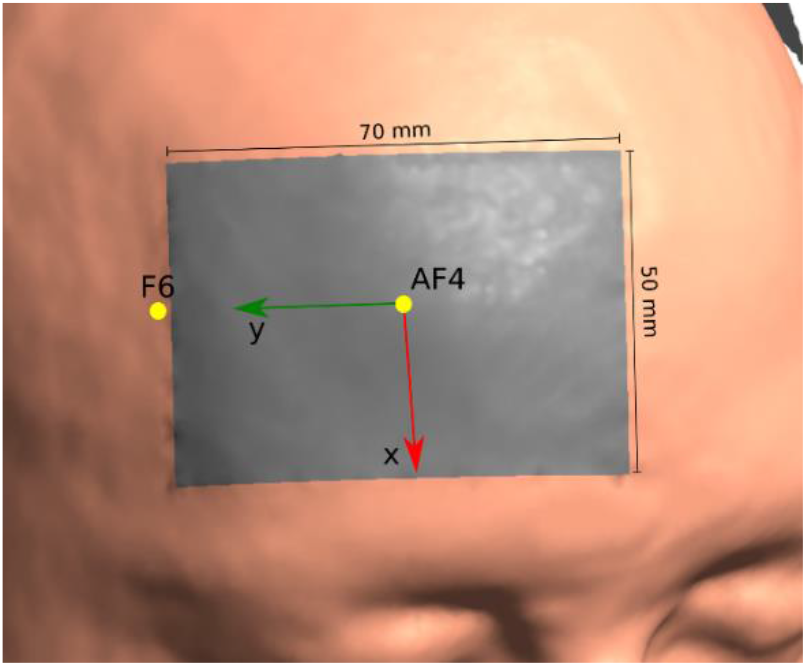
50×70 mm electrode defined with a “centre” in AF4 and a “pos_ydir “ in F6.

When the *map_to_fsavg* option is set to *true*, SimNIBS computes the electric fields in a surface located in the middle of the GM layer. This cortical surface, along with the norm, normal and tangent components of the electric field at the cortical surface and the angle between the electric field and the cortical surface can found in the *subject_overlays* folder, for both the left hemisphere (*lh*) and for the right hemisphere (*rh*) as shown in Table 5. Afterwards, these quantities are transformed into the FsAverage space. The transformed quantities can be found in the *fsavg_overlays* folder, as shown in table 5. Additionally, we have the electric field and its norm in MNI space in the *mni_volumes* folder. When using the MNI space for group analysis, it is highly recommended to use the current density “J”, or its norm “normJ” instead of the electric field “E” or its norm “normE”. This is because the current density, unlike the electric field, is continuous across tissue boundaries, and therefore more resilient to inaccuracies of the registration to the MNI template.

#### 3.2.4 Visualizing Results

We can also make use of the MATLAB library of SimNIBS to analyse the results from the simulations. Here, we are interested in the average and standard deviation of the normal component of the electric field in the cortex. The normal component, as shown in Figure 7 is the part of the electric field which is either entering or leaving the cortex.

Script 2 loads the normal field component data for each subject and calculates the mean and the standard deviation across subjects at each position of the FsAverage template. The fields are then visualized using MATLAB visualization tools. The results are shown in Figure 11. We can for example see strong current in-flow in the central gyrus, and large variations in the normal component in frontal regions.

**Script 2:**
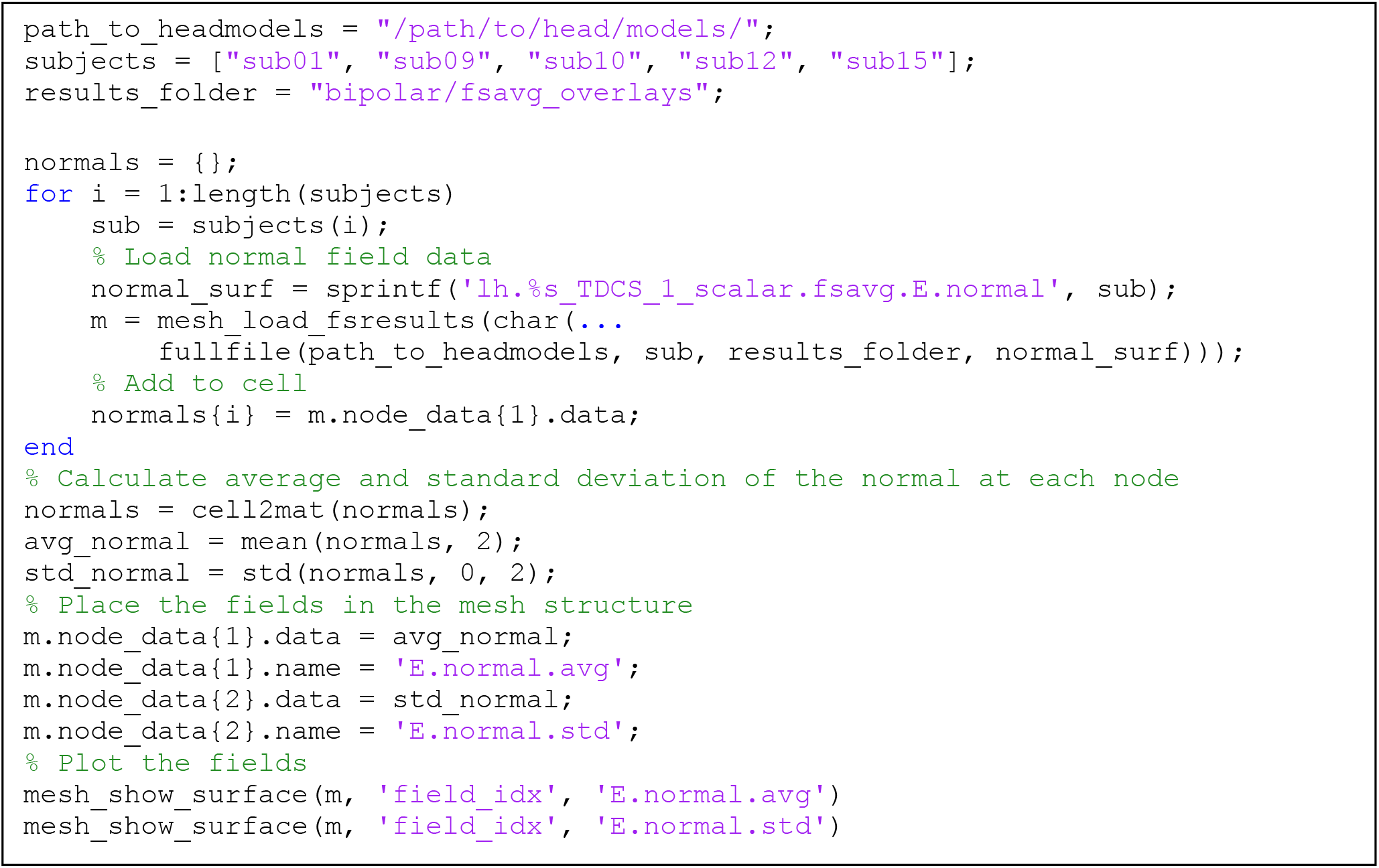
Analysis of simulation results in FSAverage space.

**Figure 11:**
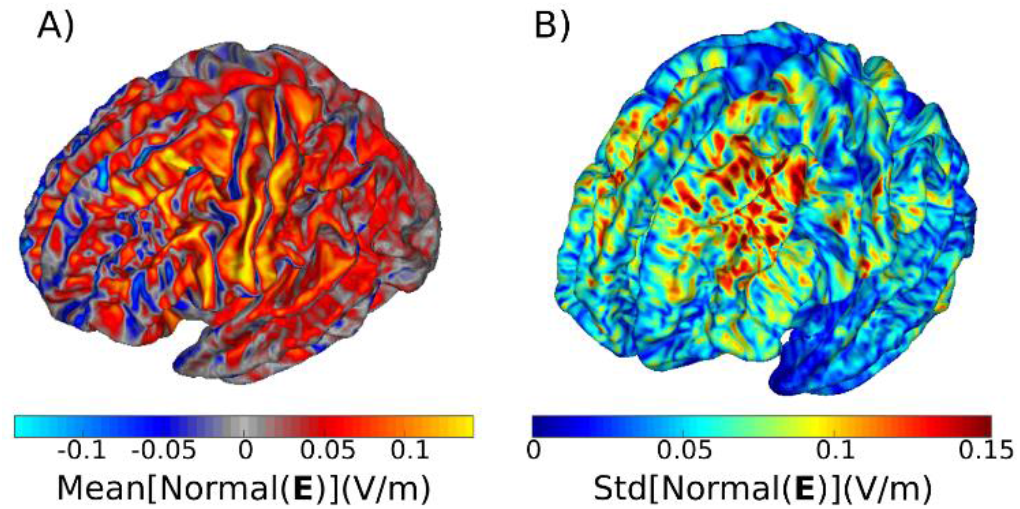
A) Mean and B) Standard deviation of the normal field component across 5 subjects. The fields were caused by tDCS with an anode over C3 and a cathode over AF4. Positive values in A) denote inflowing currents, and negative values outflowing currents.

## 4. The accuracy of automatic EEG positioning

Here, we compare EEG 10-10 positions obtained either from:

A. Transforming EEG 10-10 electrode positions defined in MNI space to the subject space using a non-linear transform, and then projecting the positions to the scalp. This is done for both **mri2mesh** and **headreco** head models.
B. Manually locating the fiducials: Left Pre-Auricular point (LPA), Right Pre-Auricular point (RPA), Nasion (Nz) and Inion (Iz) on MRI images, and afterwards calculating the EEG positions using the definitions in [22].

Calculations using method A require no user input and are automatically performed in both **mri2mesh** and **headreco** head modelling pipelines, while calculations using method B requires the user to manually select the fiducial positions.

To compare the methods A and B to position the electrodes, we calculated the EEG 10-10 positions using both ways for MR data of 17 subjects (10 females, age range: 20-35 years). The data was acquired as part of a larger study. The subjects gave written informed consent before the scan, and the study was approved by the local ethics committee of the University of Greifswald (Germany). The 17 datasets were acquired on a 3-Tesla Siemens Verio scanner (Siemens Healthcare, Erlangen, Germany) using a 32-channel head coil (T1: 1×1×1 mm^3^, TR 2300 ms, TE 900 ms, flip angle 9°, with selective water excitation for fat suppression; T2: 1×1×1 mm3, TR 12770 ms, TE 86 ms, flip angle 111°). For method B, the fiducials were manually located for each subject by a trained investigator on the T1- and T2-weighted images. The rater had no knowledge of the automatically determined positions. The fiducials Nz, Iz, LPA, and RPA were set in freeview, following the procedure described in [22] and additionally verified using the SimNIBS GUI. The subject-specific coordinates of the fiducials were extracted and these manually set positions were then compared with those calculated by the automatic algorithm in each individual.

Table 6 shows the maximal distance across all subjects between the fiducials obtained using method A and manually selected fiducials (B). We see that Nz is the most consistent fiducial, where we have the least deviation, whereas Iz is where we have the highest deviation. Also, the maximal difference in position across the two methods is ~1 cm, indicating that the method A works well to approximate the positions of the fiducials.

**Table 6:** Maximum and mean distance between the fiducial positions selected by hand and obtained from the MNI transformations across 17 subjects, for the two head modelling pipelines.

| Fiducial | mri2mesh |  | headreco |  |
| --- | --- | --- | --- | --- |
|  | Max Distance (mm) | Mean Distance +- Standard Deviation (mm) | Max Distance (mm) | Mean Distance +- Standard Deviation (mm) |
| LPA | 6.4 | 3.2 +- 1.5 | 8.7 | 5.4 +- 2.0 |
| RPA | 8.9 | 3.0 +- 1.6 | 10.6 | 5.9 +- 1.7 |
| Nz | 3.9 | 2.1 +- 1.0 | 6.0 | 3.9 +- 1.6 |
| Iz | 14.3 | 4.0 +- 3.5 | 13.2 | 5.2 +- 3.3 |

Furthermore, in Figure 12 we compare the two methods for all electrode positions in the EEG 10-10 system. The deviation in positioning each electrode was calculated as the mean of the distance between the positions obtained with either headreco or mri2mesh to the manually located fiducial positions, across all 17 subjects and for each electrode.

**Figure 12:**
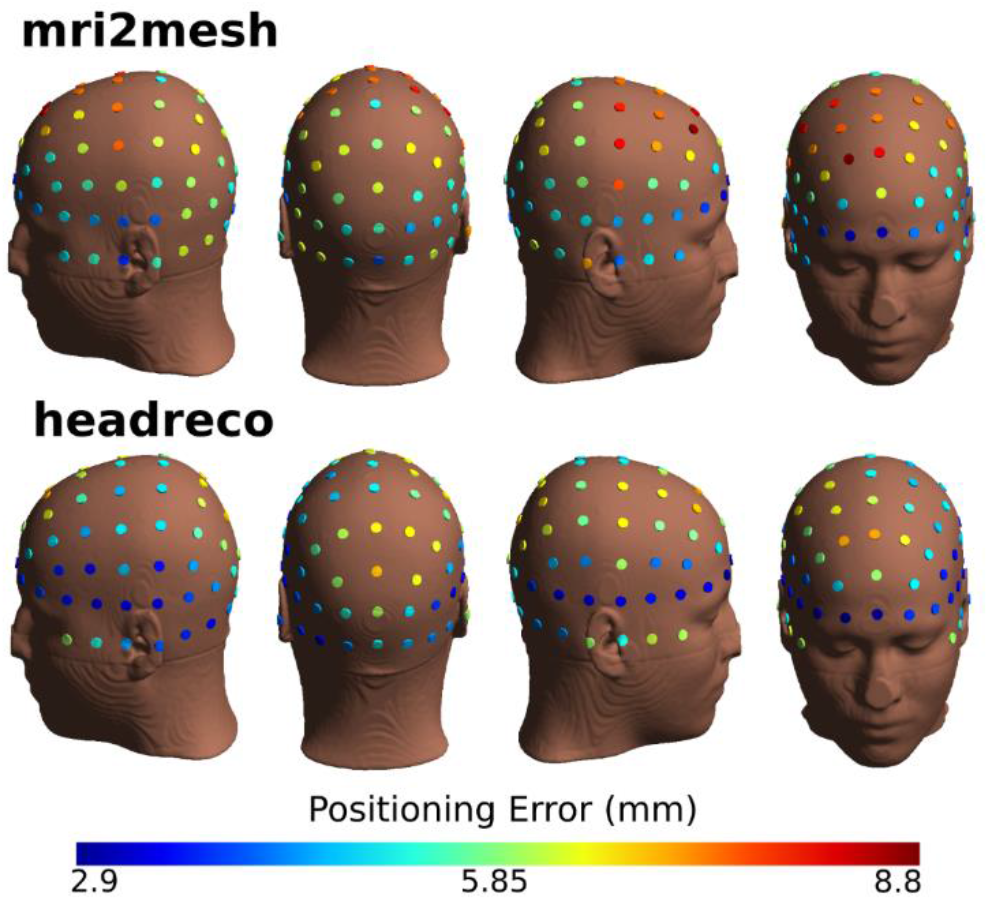
Positioning error for electrodes in the EEG 10-10 system. The error is calculated by comparing the positions calculated based on manually selected fiducials to positions calculated based on non-linear MNI transformations.

The errors for all electrodes are below 1 cm, indicating that the two algorithms for placing EEG electrodes are in agreement. We can also see that the errors in the EEG positions obtained from **headreco** are on average lower than the ones obtained from **mri2mesh**. It also seems that the anterior electrodes have less errors than the posterior electrodes. Interestingly, the location of the errors is different across the two pipelines, with **mri2mesh** being more inaccurate in superior regions and **headreco** more inaccurate in posterior regions. This might be because the way FSL (**mri2mesh**) and SPM (**headreco**) calculate non-linear MNI transformations is different. The average error across all positions was of 5.6 mm for **mri2mesh** head models and 4.9 mm for **headreco** head models indicating good accuracy.

## 5. Conclusion

We presented SimNIBS 2.1 (www.simnibs.org), a software for individualized modelling of electric fields caused by non-invasive brain stimulation. SimNIBS is free software and avaliable for all major platforms. SimNIBS does not require the installation of any additional software in order to run simulations on the example dataset. To construct head models, SimNIBS relies either on MATLAB, SPM12 and CAT12 (**headreco**) or on FSL and FreeSurfer (**mri2mesh**).

We also presented two examples of workflows in SimNIBS. In the first example, we started by using **headreco** to construct a head model. Afterwards, we used the GUI to setup a tDCS and a TMS simulation in an interactive way, and finally visualized the results. In the second example, we constructed several head models and used a MATLAB script to run simulations for each subject. Afterwards, we calculated the mean and the stardard deviation of the electric field norm across all subjects, using the FreeSurfer’s FsAverage brain template. Finally, we show results validating our automatic procedure to obtain electrode positions for the EEG 10-10 system.

SimNIBS is still being actively developed, and we expect further updates to be implemented in the future.

## Acknowledgements

Lundbeckfonden (grant Nr. R118-A11308), and NovoNordisk fonden (grant Nr. NNF14OC0011413). This project has received funding from the European Union’s Horizon 2020 research and innovation programme under grant agreement No 731827. The results and conclusions in this article present the authors’ own views, and do not reflect those of the EU Commission.

